# BABAPPAlign: A Multiple Sequence Alignment Engine with a Learned Residue-Level Scoring Function

**DOI:** 10.64898/2025.12.26.696577

**Authors:** Krishnendu Sinha

**Affiliations:** Department of Zoology, Jhargram Raj College, Jhargram–721507, West Bengal, India

**Keywords:** multiple sequence alignment, protein language models, learned scoring, affine-gap dynamic programming, BAliBASE

## Abstract

Multiple sequence alignment (MSA) underpins comparative genomics, evolutionary analysis, and structural inference. Despite extensive methodological development, most widely used alignment algorithms rely on static amino acid substitution matrices that encode global average substitution tendencies and are inherently context-agnostic. Such models are limited in their ability to capture sequence-specific evolutionary constraints, particularly for divergent or low-signal protein families.

BABAPPAlign is a multiple sequence alignment framework that replaces static substitution matrices with a learned, context-aware residue-level scoring function while preserving exact affine-gap dynamic programming. Residue compatibility is inferred using a neural scoring model, BABAPPAScore, operating on fixed contextual embeddings derived from a pretrained protein language model. Neural inference is performed entirely outside the dynamic programming recursion, ensuring that alignment optimization remains exact, deterministic, and reproducible.

Evaluation on the BAliBASE benchmark using strict family-wise paired comparisons demonstrates statistically significant improvements in sum-of-pairs and total column scores over ClustalW, MUSCLE, MAFFT, and T-Coffee across 386 reference families.

## 1 Introduction

Multiple sequence alignment (MSA) is a foundational operation in computational biology, underpinning phylogenetic reconstruction, functional motif discovery, evolutionary rate estimation, and comparative structural analysis [1, 2]. Despite decades of algorithmic development, accurate alignment of divergent protein families remains a central and unresolved challenge, particularly in regimes characterized by low sequence identity, compositional bias, and heterogeneous evolutionary constraints.

Most widely used MSA tools—including ClustalW, MUSCLE, MAFFT, and T-Coffee—are built upon variants of dynamic programming combined with progressive, iterative, or consistency-based heuristics [3, 4, 5, 6]. Although these methods differ in guide-tree construction, refinement strategies, and heuristic optimizations, they share a common reliance on static amino acid substitution matrices such as PAM and BLOSUM [7, 8]. These matrices encode global average substitution tendencies estimated across large and heterogeneous protein corpora and are therefore inherently context-agnostic. As a result, they are ill-suited to capture sequence-specific evolutionary pressures, structural constraints, or local biochemical environments, particularly in low-signal alignment regimes.

Recent advances in protein language models have demonstrated that rich contextual representations of amino acid residues can be learned directly from large collections of unaligned protein sequences [9, 10]. These models implicitly encode information about residue identity, biochemical compatibility, structural context, and long-range dependencies, suggesting that residue–residue compatibility can be inferred dynamically from sequence context rather than imposed via fixed substitution tables. While such representations have been widely adopted for structure prediction and functional annotation, their principled integration into classical sequence alignment frameworks remains limited.

Several recent alignment approaches have incorporated machine learning in various forms, including heuristic scoring adjustments, probabilistic models, or end-to-end differentiable alignment objectives. However, many such methods either replace exact dynamic programming with approximate inference or entangle representation learning, scoring, and optimization in ways that obscure the source of performance gains and complicate reproducibility. Consequently, it remains unclear whether reported improvements arise from improved residue compatibility modeling, altered optimization criteria, or heuristic shortcuts.

Here, BABAPPAlign is introduced as a multiple sequence alignment framework that isolates learning strictly to the residue-level scoring function while preserving exact affine-gap dynamic programming. In BABAPPAlign, classical substitution matrices are replaced by a learned, context-aware residue compatibility function—BABAPPAScore—operating on fixed contextual embeddings derived from a pretrained protein language model. Importantly, neural inference is performed entirely outside the dynamic programming recursion: all residue–residue or column–residue scores are computed in advance and supplied as fixed numerical inputs to the alignment algorithm. As a result, alignment optimization remains exact, interpretable at the algorithmic level, and fully reproducible.

This design occupies a distinct position in the MSA method landscape. By preserving the Needleman–Wunsch–Gotoh formulation and modifying only the scoring function, BABAPPAlign enables a direct assessment of how learned residue compatibility affects alignment accuracy, independent of heuristic alignment strategies or altered objective functions. The framework is modular by construction, allowing alternative embedding models or scoring functions to be substituted without modifying the alignment algorithm itself.

BABAPPAlign is evaluated on the BAliBASE benchmark using strict family-wise paired comparisons and nonparametric statistical analyses. All BAliBASE reference families are excluded from training, validation, and model selection, ensuring that reported performance reflects genuine generalization. Across the full benchmark and across alignment difficulty categories, BABAPPAlign demonstrates consistent and statistically robust improvements in alignment accuracy, with the largest gains observed in challenging and biologically informative regimes. These results establish learned residue-level scoring as a principled and extensible alternative to static substitution matrices, while retaining the optimality, transparency, and reproducibility of classical alignment algorithms.

### 2 Methods

### 2.1 Methodological Overview

BABAPPAlign is a multiple sequence alignment (MSA) framework that explicitly separates residue representation, residue–residue scoring, and alignment optimization. Given a set of protein sequences, contextual residue representations are first computed using a pretrained protein language model. A learned residue-level scoring function, BABAPPAScore, then converts these representations into numerical compatibility scores. These scores are supplied as fixed inputs to an exact affine-gap dynamic programming algorithm, which performs alignment optimization without heuristic approximation.

Multiple sequence alignments are constructed progressively using a deterministic profile– sequence strategy. The progressive order serves solely as a computational mechanism for assembling the alignment and is not interpreted as a phylogenetic reconstruction. An overview of the framework is shown in Figure 1.

**Figure 1.**
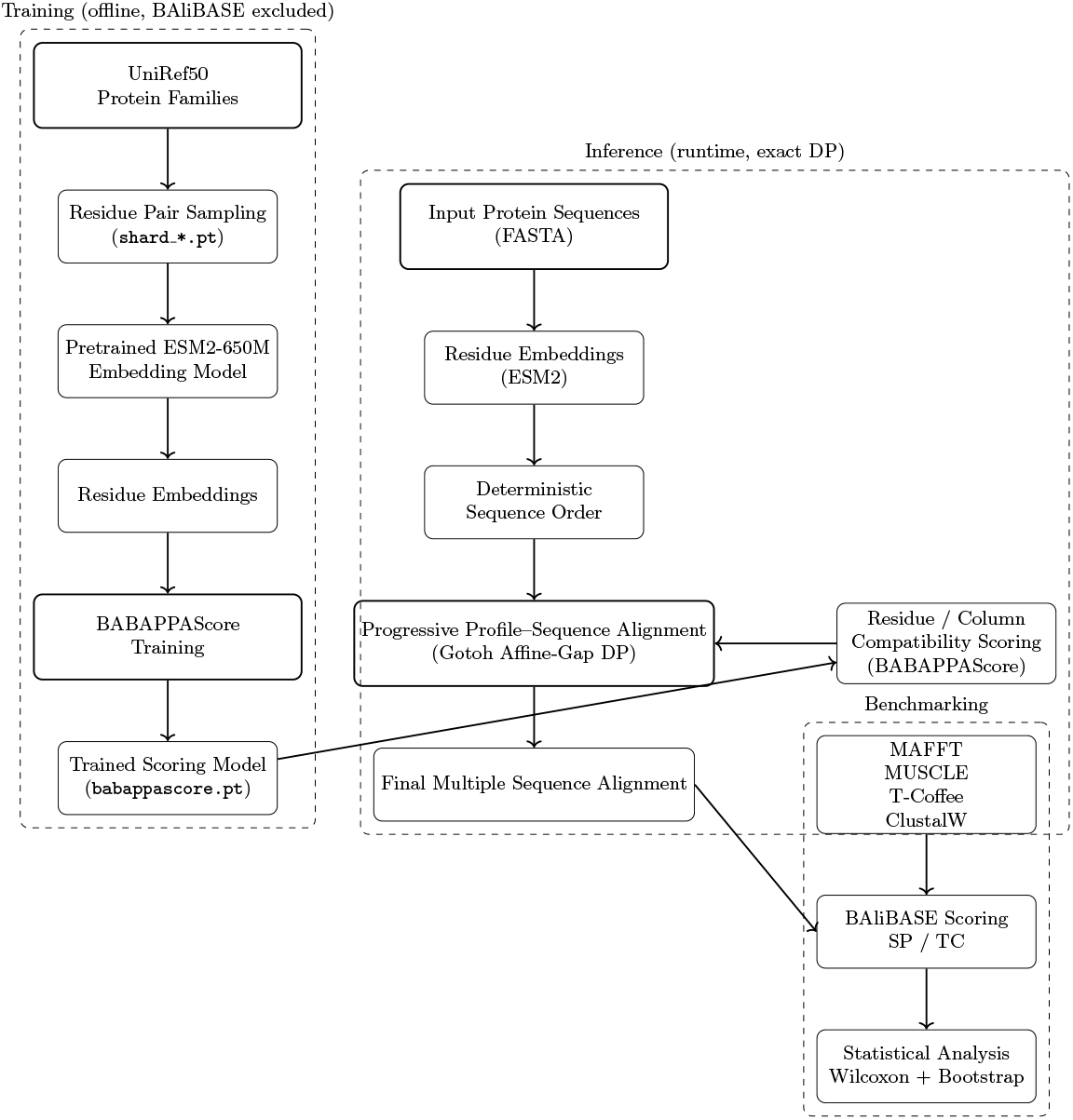
Overview of the BABAPPAlign framework. A learned residue-level scoring model (BABAPPAScore), trained offline with BAliBASE excluded, provides fixed compatibility scores that parameterize exact affine-gap dynamic programming. Multiple sequence alignment is constructed via deterministic progressive profile–sequence alignment without guide-tree inference. Neural inference is performed entirely outside the dynamic programming recursion.

### 2.2 Residue Representations from Protein Language Models

Residue-level representations were generated using a pretrained transformer-based protein language model from the ESM-2 family (`esm2_t33_650M_UR50D`). For a protein sequence

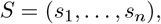

the model produces contextual embeddings

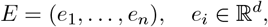

where each *e*_*i*_ encodes information about residue *s*_*i*_ conditioned on its full sequence context. Embeddings were computed in feature-extraction mode with fixed parameters, cached for reuse, and not fine-tuned, ensuring that downstream performance differences arise from the scoring model rather than representation learning.

### 2.3 BABAPPAScore: Learned Residue-Level Compatibility Function

Residue compatibility is quantified using BABAPPAScore, a learned scoring function designed to replace static amino acid substitution matrices. For two sequences *S*^(*A*)^ and *S*^(*B*)^ with embeddings *E*^(*A*)^ and *E*^(*B*)^, compatibility between residues *i* and *j* is defined as

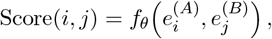

where *f*_*θ*_ is a parameterized neural function. The function is explicitly symmetric,

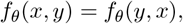

ensuring consistency under sequence reordering and compatibility with classical dynamic programming frameworks.

For each sequence–sequence or profile–sequence comparison, BABAPPA Score produces a dense numerical score matrix that directly replaces classical substitution matrices, allowing residue compatibility to depend on sequence-specific contextual information.

### 2.4 Progressive Profile–Sequence Alignment

Multiple sequence alignment is performed using a deterministic progressive profile–sequence strategy without constructing an explicit guide tree or distance matrix. Given an input ordering (*S*_1_, …, *S*_*N*_ ), alignment is initialized with *S*_1_ and extended by aligning the current profile to each subsequent sequence using exact global affine-gap dynamic programming.

Profile–sequence alignment follows the Needleman–Wunsch formulation with affine gap penalties as described by Gotoh[2]. Let *M* (*i, j*), *I*_*x*_(*i, j*), and *I*_*y*_(*i, j*) denote the match, insertion, and deletion states. Match scores are parameterized by BABAPPAScore evaluated between profile column embeddings and residue embeddings. All neural scoring is performed prior to dynamic programming; the alignment recursion itself operates exclusively on fixed numerical scores, ensuring exact and deterministic optimization.

### 2.5 Training Data and Benchmark Isolation

Training data for BABAPPAScore were derived from curated protein families obtained from UniRef50. All BAliBASE reference families were strictly excluded from training, validation, and model selection to prevent information leakage. BAliBASE datasets were reserved exclusively for benchmarking.

Protein families were processed into standardized FASTA format and organized into deterministic shards to enable scalable preprocessing and exact reproducibility.

### 2.6 Benchmarking Protocol

Benchmarking was conducted using a dedicated evaluation pipeline on the BAliBASE 3 reference dataset (RV11–RV50). BABAPPAlign was evaluated alongside MAFFT, MUSCLE, T-Coffee, and ClustalW using standard, recommended command-line settings. All tools were run on identical unaligned input sequences, with a fixed per-family timeout.

Alignment accuracy was assessed using a single BAliBASE scoring implementation to compute sum-of-pairs (SP) and total column (TC) scores. Benchmarking was performed serially to eliminate nondeterminism. Execution logs, runtimes, tool versions, and failure modes were recorded. Families with failed alignments for any method were excluded from paired comparisons.

### 2.7 Statistical Analysis and Reproducibility

All comparisons were conducted as paired, family-wise analyses. Differences in SP, TC, and runtime were evaluated using the Wilcoxon signed-rank test. Effect sizes were quantified using Cliff’s delta, and 95% confidence intervals were estimated via nonparametric bootstrap resampling. False discovery rate correction was applied using the Benjamini–Hochberg procedure.

BABAPPAlign is deterministic given fixed inputs, model weights, and parameters. Residue embeddings were cached, scoring models were versioned, and alignment parameters were held constant. Experiments were conducted on Linux systems; GPU acceleration affected performance only, not correctness. Under identical conditions, BABAPPAlign produces identical alignments across runs.

## 3 Results

### 3.1 Overall BAliBASE Benchmark

Overall alignment performance was evaluated on 386 BAliBASE reference families using paired family-wise comparisons. For each family, BABAPPAlign and all comparator methods (ClustalW, MUSCLE, MAFFT, and T-Coffee) were run on identical input sequences, and differences in alignment quality were computed on a per-family basis. This paired design ensures that all reported effects reflect method-specific performance rather than variation in dataset composition. Across the full benchmark, BABAPPAlign consistently achieved higher sum-of-pairs (SP) and total column (TC) scores than all comparator methods (Table 1). Median improvements in SP score ranged from 0.025 (relative to MAFFT) to 0.067 (relative to ClustalW), with corresponding Cliff’s delta values indicating medium to large effect sizes. Improvements in TC score followed a similar trend, though with smaller magnitudes, reflecting the more conservative nature of the TC metric.

**Table 1.** Overall BAliBASE benchmarking statistics comparing BABAPPAlign against established MSA tools. All values are derived from paired comparisons across 386 BAliBASE families.

| Metric | Comparator | $\Delta$ Median | 95% CI | Cliff's $\delta$ | FDR $p$ |
| --- | --- | --- | --- | --- | --- |
| SP | ClustalW | 0.0669 | [0.0547, 0.0837] | 0.751 | $2.7 \times 10^{-48}$ |
| SP | MAFFT | 0.0252 | [0.0190, 0.0306] | 0.523 | $3.2 \times 10^{-29}$ |
| SP | MUSCLE | 0.0512 | [0.0419, 0.0664] | 0.626 | $8.7 \times 10^{-40}$ |
| SP | T-Coffee | 0.0642 | [0.0536, 0.0782] | 0.668 | $5.2 \times 10^{-46}$ |
| TC | ClustalW | 0.0665 | [0.0534, 0.0785] | 0.491 | $2.1 \times 10^{-27}$ |
| TC | MAFFT | 0.0255 | [0.0187, 0.0316] | 0.393 | $8.7 \times 10^{-18}$ |
| TC | MUSCLE | 0.0183 | [0.0115, 0.0236] | 0.279 | $4.2 \times 10^{-10}$ |
| TC | T-Coffee | 0.0382 | [0.0292, 0.0470] | 0.495 | $1.2 \times 10^{-22}$ |

All observed improvements were highly statistically significant under paired Wilcoxon signedrank testing after false discovery rate correction. Bootstrap confidence intervals confirmed that the observed median differences were well separated from zero for all tool comparisons. Runtime comparisons indicate that BABAPPAlign is computationally more expensive than heuristic aligners, reflecting the cost of learned scoring and exact affine-gap dynamic programming; however, runtime differences were highly consistent across families (Table 1).

The distribution of paired score differences is visualized in Figure 2. No statistical annotations are included in the figure; all inferential statistics and effect sizes are reported explicitly in Table 1.

**Figure 2.**
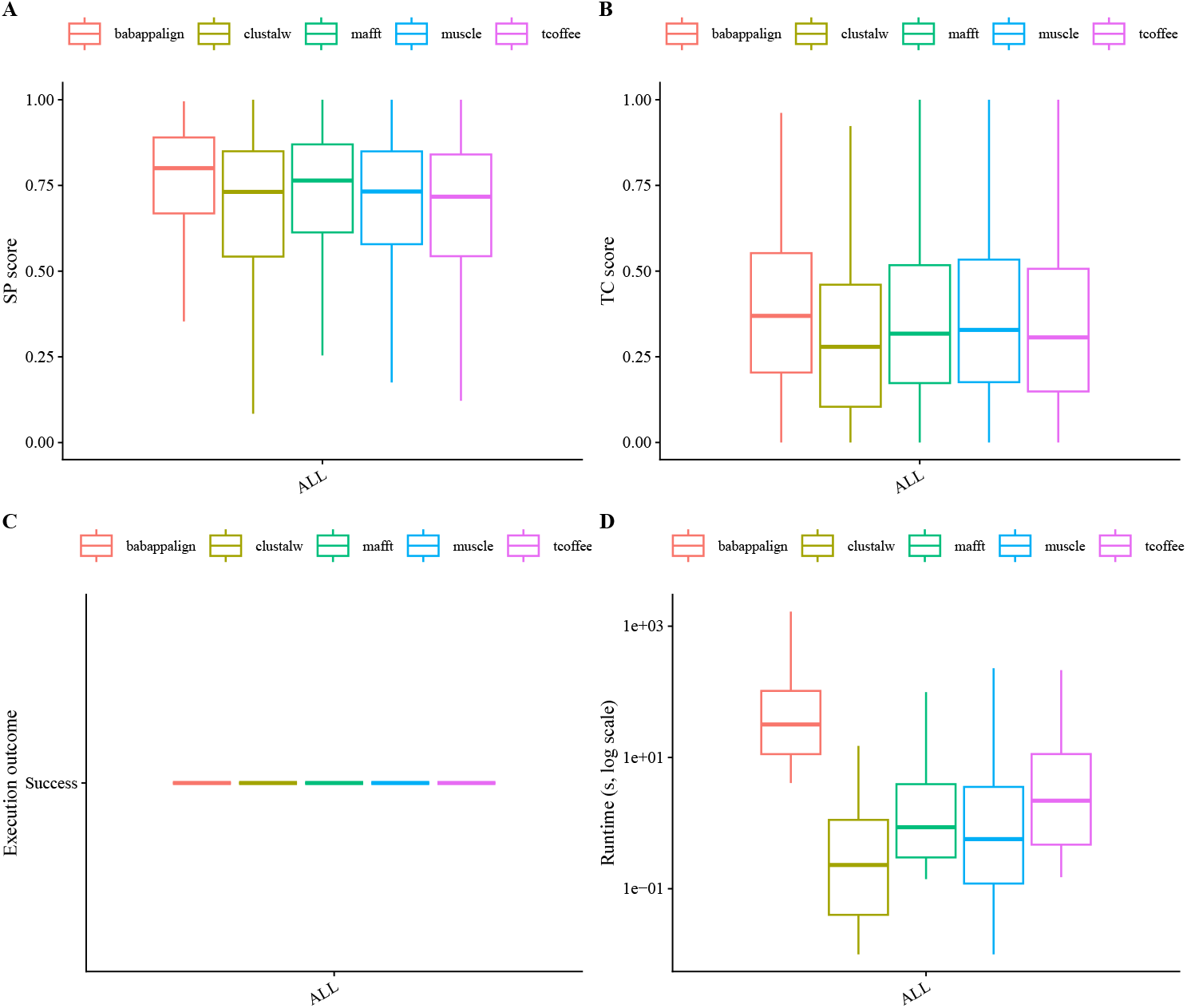
Overall BAliBASE benchmarking results across 386 reference families. Boxplots show paired family-wise differences (BABAPPAlign minus comparator) in sum-of-pairs (SP) score, total column (TC) score, and runtime. Positive values indicate superior performance of BABAPPAlign. Statistical tests, effect sizes, and confidence intervals are reported in Table 1.

### 3.2 Performance Across Alignment Difficulty Categories

To examine how alignment difficulty modulates relative method performance, BAliBASE families were stratified according to their reference variability (RV) categories, which reflect increasing uncertainty and structural heterogeneity in the reference alignment, ranging from low variability (RV11) to highly variable and ambiguous cases (RV50). Within each RV category, paired familywise comparisons between BABAPPAlign and classical aligners were conducted independently using the same nonparametric statistical framework applied in the overall benchmark.

Across RV20 and RV30 categories, which represent intermediate to high reference variability and are widely regarded as among the most informative and challenging BAliBASE subsets, BABAPPAlign exhibited consistent and substantial improvements in both SP and TC scores, accompanied by large effect sizes (Table 2). These regimes are characterized by reduced global sequence identity, frequent insertions and deletions, and locally conserved motifs embedded within divergent backgrounds. In such settings, learned residue-level scoring confers a clear advantage by adapting dynamically to sequence-specific and contextual signals that are poorly captured by static substitution matrices.

**Table 2.**
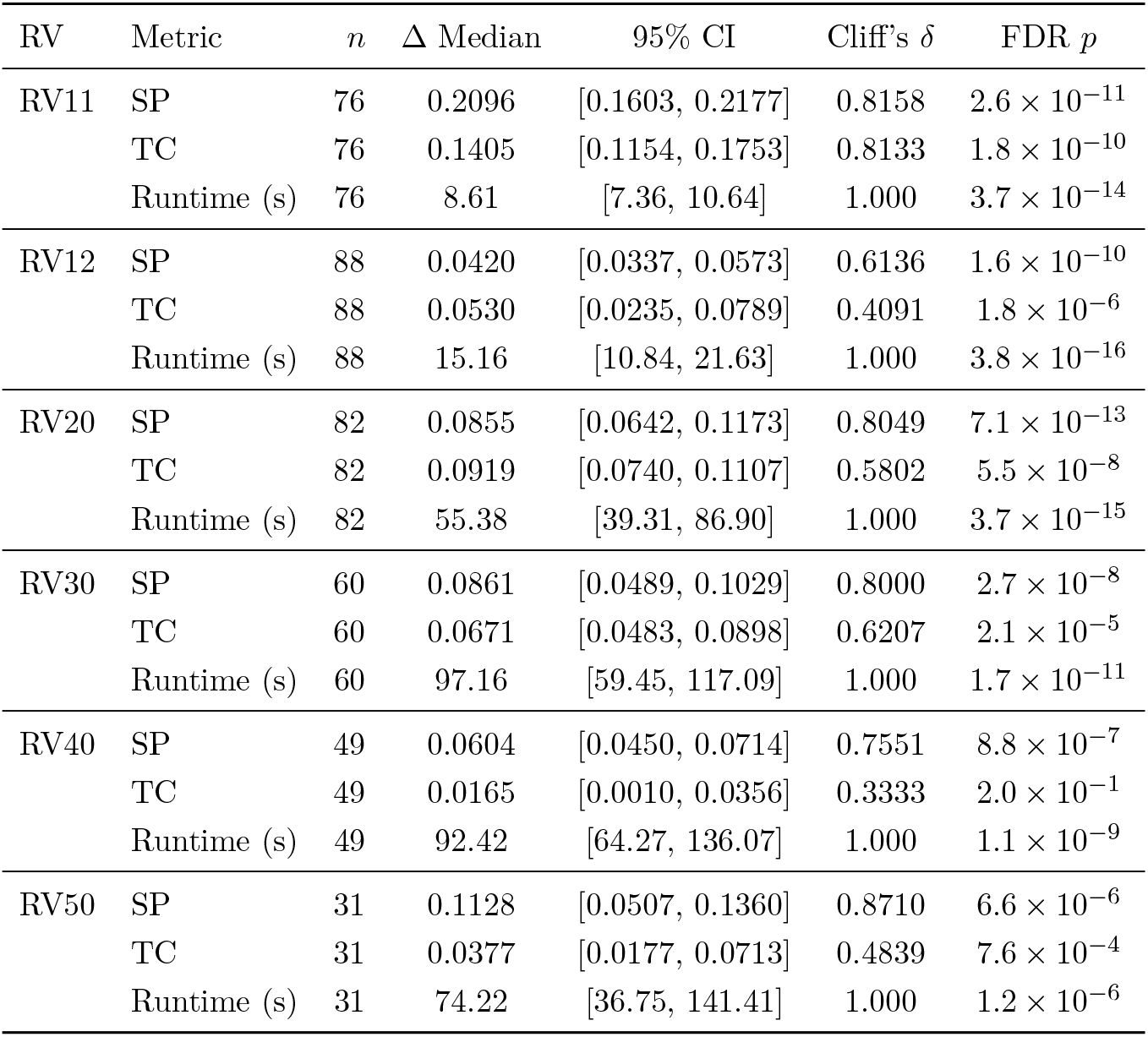
RV-wise benchmarking results on BAliBASE stratified by reference difficulty category. For each RV group, paired family-wise comparisons are reported as median differences (BABAPPAlign minus comparator), 95% bootstrap confidence intervals, Cliff’s delta (effect size), and FDR-adjusted *p* values. Positive Δ indicates superior performance of BABAPPAlign.

| RV | Metric | $n$ | $\Delta$ Median | 95% CI | Cliff’s $\delta$ | FDR $p$ |
| --- | --- | --- | --- | --- | --- | --- |
| RV11 | SP | 76 | 0.2096 | [0.1603, 0.2177] | 0.8158 | $2.6 \times 10^{-11}$ |
| | TC | 76 | 0.1405 | [0.1154, 0.1753] | 0.8133 | $1.8 \times 10^{-10}$ |
| | Runtime (s) | 76 | 8.61 | [7.36, 10.64] | 1.000 | $3.7 \times 10^{-14}$ |
| RV12 | SP | 88 | 0.0420 | [0.0337, 0.0573] | 0.6136 | $1.6 \times 10^{-10}$ |
| | TC | 88 | 0.0530 | [0.0235, 0.0789] | 0.4091 | $1.8 \times 10^{-6}$ |
| | Runtime (s) | 88 | 15.16 | [10.84, 21.63] | 1.000 | $3.8 \times 10^{-16}$ |
| RV20 | SP | 82 | 0.0855 | [0.0642, 0.1173] | 0.8049 | $7.1 \times 10^{-13}$ |
| | TC | 82 | 0.0919 | [0.0740, 0.1107] | 0.5802 | $5.5 \times 10^{-8}$ |
| | Runtime (s) | 82 | 55.38 | [39.31, 86.90] | 1.000 | $3.7 \times 10^{-15}$ |
| RV30 | SP | 60 | 0.0861 | [0.0489, 0.1029] | 0.8000 | $2.7 \times 10^{-8}$ |
| | TC | 60 | 0.0671 | [0.0483, 0.0898] | 0.6207 | $2.1 \times 10^{-5}$ |
| | Runtime (s) | 60 | 97.16 | [59.45, 117.09] | 1.000 | $1.7 \times 10^{-11}$ |
| RV40 | SP | 49 | 0.0604 | [0.0450, 0.0714] | 0.7551 | $8.8 \times 10^{-7}$ |
| | TC | 49 | 0.0165 | [0.0010, 0.0356] | 0.3333 | $2.0 \times 10^{-1}$ |
| | Runtime (s) | 49 | 92.42 | [64.27, 136.07] | 1.000 | $1.1 \times 10^{-9}$ |
| RV50 | SP | 31 | 0.1128 | [0.0507, 0.1360] | 0.8710 | $6.6 \times 10^{-6}$ |
| | TC | 31 | 0.0377 | [0.0177, 0.0713] | 0.4839 | $7.6 \times 10^{-4}$ |
| | Runtime (s) | 31 | 74.22 | [36.75, 141.41] | 1.000 | $1.2 \times 10^{-6}$ |

Notably, statistically significant gains were also observed in RV11, despite its low reference variability. This indicates that the learned scoring model does not merely compensate for extreme divergence but can also refine alignments in well-defined reference regimes, suggesting improved sensitivity to subtle residue-level compatibility even when structural agreement is strong.

In contrast, performance differences in RV40 and RV50 were generally smaller and, for the TC metric, occasionally not statistically significant. These categories are characterized by high reference variability and intrinsic ambiguity in column definitions, which increases variance in TC-based evaluation and limits the sensitivity of column-level metrics to detect incremental improvements. Importantly, no systematic degradation in alignment quality was observed for BABAPPAlign in any RV category, indicating that performance gains in more informative regimes are not offset by losses in highly variable cases.

Figure 3 summarizes the distributions of paired score differences across RV classes. As in the overall benchmark, figures are presented without statistical annotations; all confidence intervals, effect sizes, and multiple-testing–corrected significance values are reported explicitly in Table 2. Collectively, these results demonstrate that the performance advantages of BABAPPAlign are not driven by trivial or highly conserved alignments, but are most pronounced in biologically informative regimes where classical scoring models are intrinsically limited and alignment decisions are most consequential.

**Figure 3.**
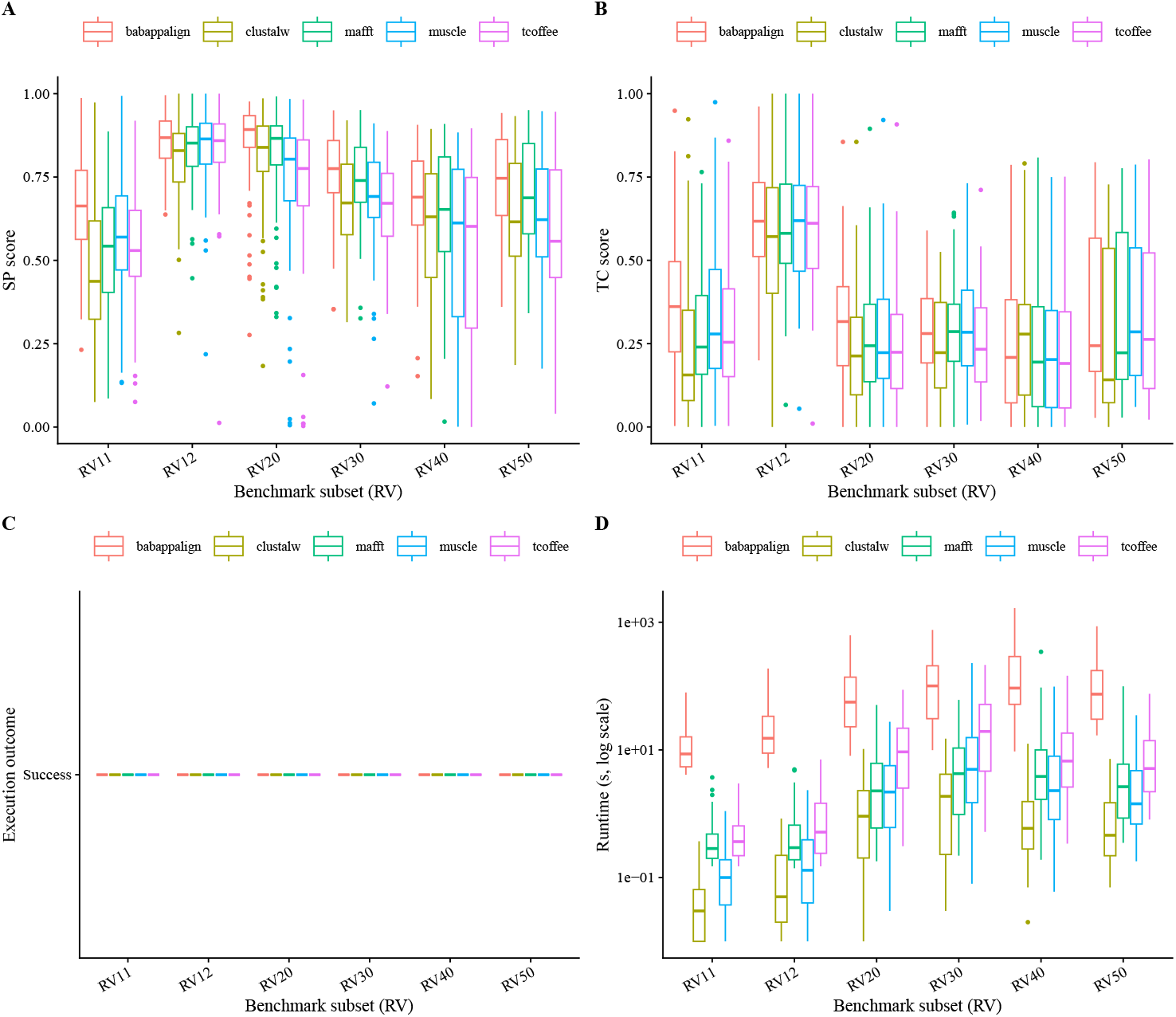
BAliBASE benchmarking results stratified by reference variability (RV) category. Boxplots show paired family-wise differences (BABAPPAlign minus comparator) in SP score, TC score, and runtime within each RV class. Positive values indicate higher alignment accuracy or increased runtime for BABAPPAlign. Statistical analyses are reported in Table 2.

## 4 Discussion and Conclusion

The results presented here demonstrate that learned residue-level scoring can be integrated into classical affine-gap dynamic programming to yield consistent and statistically robust improvements in multiple sequence alignment accuracy. Unlike approaches that modify alignment optimization itself or rely on heuristic inference, BABAPPAlign isolates learning strictly to the residue compatibility model while preserving exact dynamic programming. This separation makes it possible to attribute performance gains unambiguously to improved scoring rather than altered optimization criteria, heuristic shortcuts, or stochastic training effects.

A central implication of this work is that static substitution matrices constitute a fundamental bottleneck for alignment accuracy in diverse evolutionary regimes. By construction, matrices such as PAM and BLOSUM encode global average substitution tendencies and cannot adapt to sequence-specific context, local biochemical environments, or heterogeneous evolutionary constraints. In contrast, BABAPPAScore leverages contextual residue embeddings to infer compatibility dynamically, enabling alignment decisions to respond to both local and long-range sequence information. The consistent improvements observed across BAliBASE families, particularly in intermediate and high-difficulty categories, indicate that context-aware scoring provides actionable signal precisely where classical models are most limited.

Importantly, the observed performance gains are not confined to highly divergent or ambiguous alignments. Statistically significant improvements are also detected in low-variability reference categories, suggesting that learned scoring can refine residue correspondence even in regimes where structural agreement is strong. At the same time, no systematic degradation in alignment quality is observed in the most variable BAliBASE categories, indicating that the learned scoring model does not introduce instability or overfitting in intrinsically ambiguous alignment regimes. Together, these results support the conclusion that learned residue-level scoring improves alignment sensitivity without sacrificing robustness.

BABAPPAlign occupies a distinct methodological position within the landscape of alignment algorithms. By retaining the Needleman–Wunsch–Gotoh formulation and modifying only the scoring function, the framework preserves algorithmic interpretability at the alignment level: explicit score matrices, well-defined gap penalties, and exact optimization. This contrasts with end-to-end neural alignment models, where scoring and optimization are entangled and the source of accuracy improvements can be difficult to disentangle. The modular design of BABAPPAlign further ensures that advances in protein representation learning can be incorporated directly, without reengineering the alignment algorithm itself.

Several limitations should be acknowledged. First, BABAPPAlign employs a deterministic progressive profile–sequence alignment order, which, like all progressive strategies, may introduce order sensitivity. While this design choice isolates the effect of learned scoring and ensures reproducibility, future extensions could incorporate guide-tree–based ordering, iterative refinement, or profile–profile alignment without altering the core scoring methodology. Second, runtime is higher than that of heuristic aligners, reflecting the cost of context-aware scoring and exact affine-gap optimization. However, runtime variance across families is low and predictable, and residue embeddings are computed once and cached, ensuring that alignment cost scales independently of model training. Finally, the current implementation focuses on protein sequence alignment; extension to nucleotide sequences or mixed alphabets would require retraining or redesign of the scoring model.

Looking forward, the separation of representation, scoring, and optimization introduced here opens several promising directions. Alternative protein language models, lightweight embedding architectures, or domain-specific representations can be substituted directly into the framework. Learned scoring could be extended to profile–profile alignment, enabling fully learned profile compatibility while retaining exact optimization. More broadly, the results suggest that learned, context-aware scoring functions may serve as a general replacement for static substitution matrices across a wide range of alignment and comparative sequence analysis tasks.

In conclusion, BABAPPAlign establishes learned residue-level scoring as a principled and extensible alternative to static substitution matrices within classical dynamic programming frameworks. By preserving exact affine-gap optimization and isolating learning to the scoring function, the approach combines the strengths of modern protein representation learning with the transparency, reproducibility, and theoretical guarantees of classical alignment algorithms. This work demonstrates that substantial improvements in multiple sequence alignment accuracy can be achieved without abandoning foundational algorithmic principles, providing a clear path for future methodological advances.

## 5 Software Availability and Reproducibility

BABAPPAlign is released as open-source software and distributed through the Bioconda package repository, enabling installation via standard Conda environments on Linux systems using conda install -c bioconda babappalign. The Bioconda package provides a CPU-compatible implementation and installs all required dependencies automatically; GPU acceleration is optional and affects performance only, not correctness. BABAPPAlign requires an external pretrained neural residue-level scoring model (BABAPPAScore), which is distributed separately via Zenodo and is not downloaded implicitly. Versioned Zenodo DOIs are provided to ensure exact reproducibility of all reported results. All experiments are fully reproducible: BABAPPAlign is deterministic given fixed inputs, model weights, and parameters, and all benchmarks were performed using identical input FASTA files, scoring model versions, and alignment parameters across methods, with BAliBASE reference families excluded from training and used exclusively for benchmarking. The source code, pretrained scoring models, and benchmarking scripts are publicly available at https://github.com/sinhakrishnendu/BABAPPAlign, with archived model releases accessible via persistent Zenodo identifiers at https://doi.org/10.5281/zenodo.18053200.

## 8 Acknowledgment

I am deeply grateful to my young son, whose boundless curiosity and love for butterflies inspired the name “BABAPPA.” In his imaginative vocabulary, “BABAPPA” refers to a butterfly, and it was this creativity that ultimately gave the software its name. I also thank my wife, Dr. Nabanita Ghosh (Maulana Azad College, Kolkata, India), for in-depth academic discussions, critical insights, and thoughtful suggestions that substantially improved the conceptual clarity of this work. Finally, I acknowledge the developers and maintainers of the open-source software tools and biological databases used by BABAPPAlign, whose contributions made this work possible.

